# Identifying SARS-CoV-2 Antiviral Compounds by Screening for Small Molecule Inhibitors of Nsp3 Papain-like Protease

**DOI:** 10.1101/2021.04.07.438804

**Authors:** Chew Theng Lim, Kang Wei Tan, Mary Wu, Rachel Ulferts, Lee A. Armstrong, Eiko Ozono, Lucy S. Drury, Jennifer C. Milligan, Theresa U. Zeisner, Jingkun Zeng, Florian Weissmann, Berta Canal, Ganka Bineva-Todd, Michael Howell, Nicola O’Reilly, Rupert Beale, Yogesh Kulathu, Karim Labib, John F.X Diffley

## Abstract

The COVID-19 pandemic has emerged as the biggest life-threatening disease of this century. Whilst vaccination should provide a long-term solution, this is pitted against the constant threat of mutations in the virus rendering the current vaccines less effective. Consequently, small molecule antiviral agents would be extremely useful to complement the vaccination program. The causative agent of COVID-19 is a novel coronavirus, SARS-CoV-2, which encodes at least nine enzymatic activities that all have drug targeting potential. The papain-like protease (PLpro) contained in the nsp3 protein generates viral non-structural proteins from a polyprotein precursor, and cleaves ubiquitin and ISG protein conjugates. Here we describe the expression and purification of PLpro. We developed a protease assay that was used to screen a custom chemical library from which we identified Dihydrotanshinone I and Ro 08-2750 as compounds that inhibit PLpro in protease and isopeptidase assays and also inhibit viral replication in cell culture-based assays.

## Introduction

By early January 2021, COVID-19 infections and deaths across 191 countries/regions were reported to be 87 million and 1.9 million respectively [1]. These numbers occurred in just over one year since the first COVID-19 outbreak was reported in December 2019. COVID-19 is a highly contagious disease with a human-to-human mode of transmission [2]. Accelerated by travellers, by the end of January 2020, the World Health Organisation (WHO) declared the disease as a “public health emergency of international concern” and the first European COVID-19 case was reported in France [3].

The causative agent of COVID-19 was identified as a novel coronavirus, Severe Acute Respiratory Syndrome coronavirus 2 (SARS-CoV-2). The SARS-CoV-2 genome was first sequenced and published in early January 2020 [4]. COVID-19 falls within the subgenus *Sarvecovirus*, genus *Betacoronavirus*, and is closely related (88% nucleotide sequence identity) to a bat-derived SARS-like coronavirus, and to the virus causing SARS (SARS-CoV-1; 79%), and more distantly related to MERS-CoV (50%) the causative agent of MERS (Middle East Respiratory Syndrome) [5].

There is currently no cure for COVID-19. In June 2020, Dexamethasone, an anti-inflammatory drug, was approved by the UK government for COVID-19 patients in National Health Service (NHS) hospitals [6]. In August 2020, remdesivir was approved by the Food and Drug Administration (FDA) in the United States [7]; this is currently the only small molecule antiviral drug approved for use against COVID-19. In addition, Casirivimab and Imdevimab, monoclonal antibody drugs, were approved under Emergency Use Authorization (EUA) by the FDA in November 2020 [8]. The other focus of treatment is prevention via vaccination. Multiple mRNA-based and virus-based vaccines have been rolled out across the world with similar overall safety and effectiveness [9–11]. However, the emergence of a new variant of SARS-CoV-2 in the UK in September 2020, has raised great concern since it is 70% more transmissible [12]. Additional variants arising across the world may render the existing vaccines less efficient. Given the uncertainty of the progression of the virus and the time frame needed to vaccinate the global community, it is crucial to search for drugs to provide treatment for COVID-19 patients.

The genome of COVID-19 is arranged as shown in Figure 1A [4]. Once the virus has entered a cell, two open reading frames, ORF 1a and ORF 1ab are translated. ORF1ab is generated from an internal ribosomal frame shift. This produces polypeptides that are processed by two viral proteases to produce 16 <u>n</u>on-<u>s</u>tructural <u>p</u>roteins (nsp) 1 to 16. The main protease is the 3C-like protease (3CLpro), corresponding to nsp5. 3CLpro cleaves in between each of the nsp 4-16 proteins. The second protease, Papain-like protease (PLpro), is contained in a small domain of the nsp3 protein; it cleaves after LXGG motifs between each of the remaining nsp1-3 [13]. Interestingly, PLpro is also a deubiquitylase (DUB) and a delSGylase, cleaving after the diglycine residues of ubiquitin (Ub) and the UBL protein ISG15 (interferon-induced gene 15), respectively, and thus has roles outside polyprotein cleavage [14, 15]. In this paper we describe our search for PLpro inhibitors.

**Figure 1.**
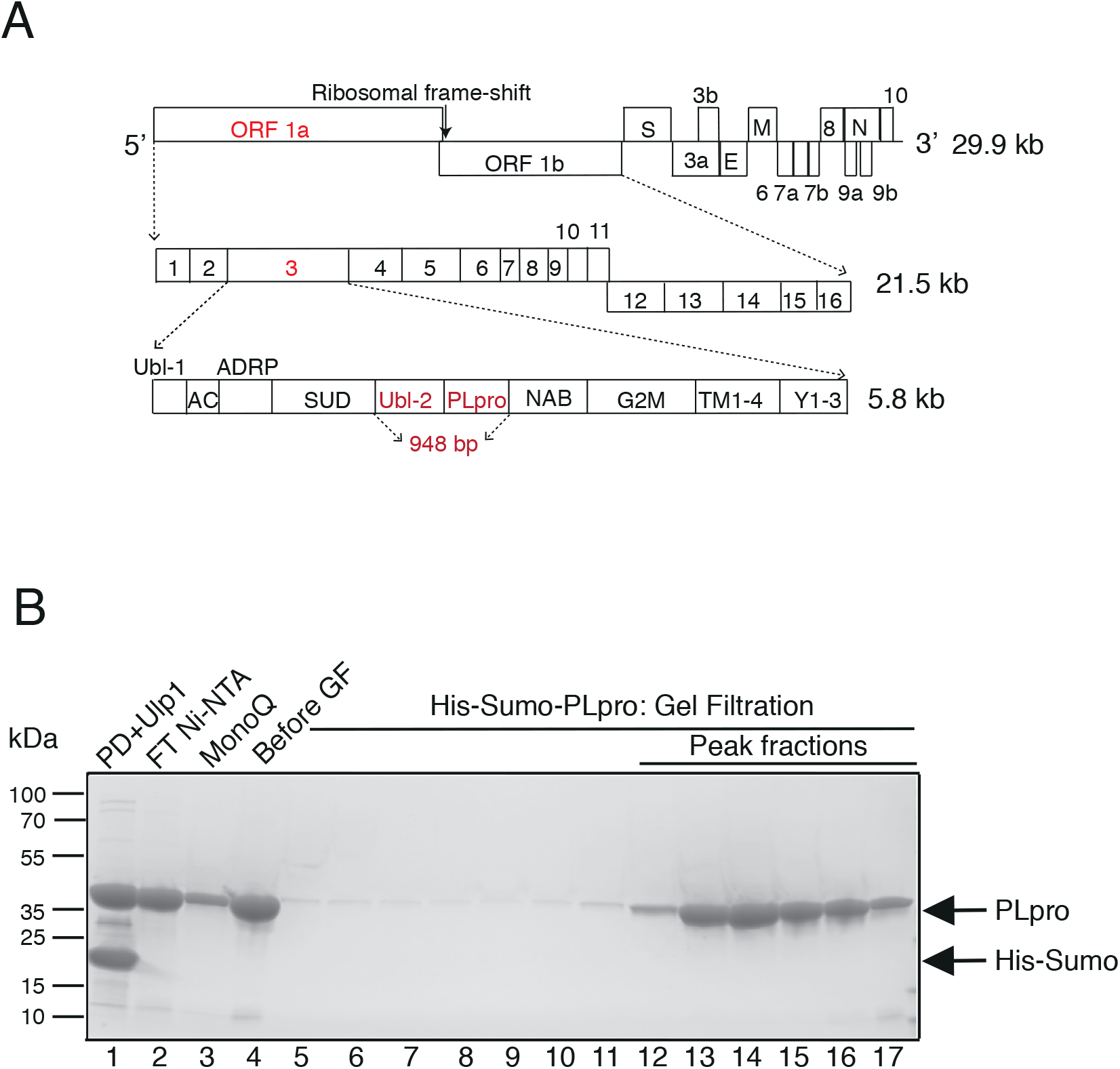
Design, expression and purification of PLpro enzyme. A. Schematic of the COVID-19 genome showing the 29.9 kb single-strand (+) RNA. ORF 1ab is 21.5 kb and codes for a polyprotein which after processing, produces 16 proteins named nsp1-16. The nsp3 protein contains the catalytic core PLpro enzyme used in this study which comprises the Ubl2 and PLpro domains (highlighted in red). The ‘948 bp’ highlighted in red indicates the sequence that was the basis of the bacterial and insect cell expression constructs used for protein purification. B. Purification of His-Sumo-PLpro. Lane 1, Digestion of the pulldown (PD) with Ulp1; lane 2, flow-through (FT) of 2^nd^ Ni-NTA; lane 3, FT from the MonoQ; lane 4, concentrated from lane 3 before applying to a gel filtration column; lanes 13-16 are the gel filtration fractions containing PLpro monomer from that were pooled for further assay.

## Results

### PLpro expression and purification

Nsp3 is the largest non-structural protein (1945 amino acids). It contains multiple domains which are arranged in the following order: ubiquitin-like (Ubl-1), acidic-domain (AC domain), ADP-ribose-1”-phosphatase (ADRP) /macro/x-domain, SARS unique Domain (SUD), Ubl-2, PLpro domain, nucleic acid-binding domain (NAB), marker domain (G2M), double-pass transmembrane domains (TM 1-2 and TM 3-4), and the Y domain (subdomains Y1-3) [14, 16] (Figure 1A). For expression and purification, we selected the region from 1564-1878 amino acids that has been previously described to produce a truncated nsp3 which encompasses the Ubl-2 and PLpro main domains [4, 14].

We expressed PLpro in both bacterial and insect cell systems to determine which versions of the purified protein retained the most enzymatic activity. For bacterial expression, PLpro was tagged with either His-Sumo or His-TEV at its N-terminus. After protein pulldown from bacterial lysate, the His-Sumo and His-TEV affinity tags were removed by Ulp1 and TEV proteases respectively (Figure 1B and Supplementary Figure S1A). For expression in insect cells, the protein was tagged with Flag-His at the N-terminus and the tag remained on the final protein (Supplementary Figure 1SB). The activities of all 3 proteins were compared as described below.

### PLpro protease activity

We used a quenched Förster (fluorescence) resonance energy transfer (FRET) technique to monitor the protease activity [17]. 2-aminobenzoyl (Abz) and a nitro-L-tyrosine (Y-(3-NO_2_)R), were added to opposite ends of a small synthetic peptide that contained the cleavage sequence recognized by PLpro. This generates a fluorescence-quenching pair (Figure 2A) in which emission from Abz is absorbed by the neighbouring Y-(3-NO_2_)R; cleavage is detected as an increase in apparent emission from Abz at 420 nM. Two cleavage sequences were initially selected which corresponded to the ten amino acids between nsp1/2 and nsp2/3, designated Pro1 and Pro2, respectively. To test for cleavage of the substrates, bacterial His-TEV-PLpro was added to the substrate in the assay buffer. Over the 20 minutes of incubation, increasing fluorescence signal was observed for substrate Pro2 but not for Pro1 (Supplementary Figure S2A). Consistent with this, it has been recently shown that full length nsp3 protein but not the isolated PLpro domain can cleave between nsp1/2 [18]. Although cleavage of Pro2 occurred, the rate was relatively slow making it less suitable for screening. We therefore designed a third substrate based on Pro2, called Pro3, which lengthened the recognition sequence peptide from 10 to 12 amino acids [19] (Figure 2A). This modification resulted in a more rapid reaction (Supplementary Figure S2B) and Pro3 was, therefore, used as substrate in subsequent experiments.

**Figure 2.**
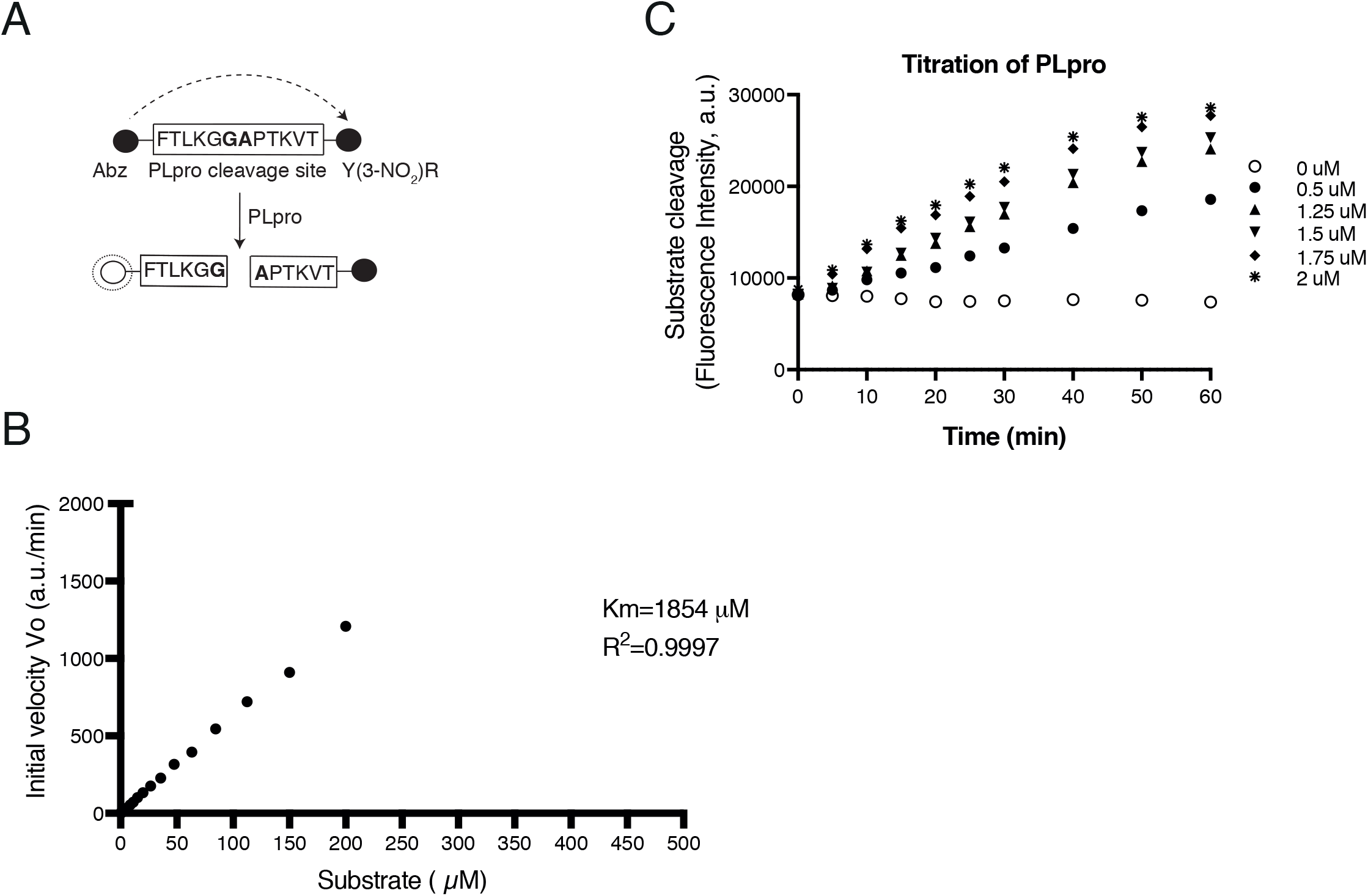
Enzyme assay design and enzyme characteristic. A. (i) Schematic of the Pro3 peptide designed for used in the FRET assay. The synthetic peptide contains 12 amino acids from the nsp2/3 junction in the natural polypeptide, in the centre of this is the FTLKGG//APTKVT sequence recognized by PLpro which cleaves between G and A. During synthesis the peptide had the fluorescent Anthranilate (2-aminobenzoyl-Abz) tag added to the N terminus and the quencher nitro-L-tyrosine (Y(3-NO_2_)R) fused to the C terminus. (ii) Organization in 384 well plate. B. Determination of the enzyme kinetics. The initial velocity of substrate hydrolysis over the titration of substrate is plotted. The Km were calculated using Michaelis-Menten equation. Data was collected from three replicates. C. Protease activity of PLpro. Titration of the purified enzyme (0.5-2 μM) incubated with the substrate. Fluorescent intensity was measured for 1 hour. Data was collected from three replicates.

The bacterial His-TEV-PLpro (tag removed) was more active than insect cell Flag-His-PLpro at the same enzyme concentration (Figure S2D). The two bacterial expression proteins, His-TEV and His-Sumo-PLpro (tags removed) showed similar activity (Figure S2C). Since the His-Sumo PLpro is an intact protein with no extra amino acids (after tag removal) and also produced a higher yield after purification, this version of PLpro was used for all of the remaining experiments.

By assaying PLpro cleavage activity across a wide range of substrate concentrations, we found that the Michaelis constant (Km) for PLpro was quite high (1854 μM) (Figure 2B). Cleavage increased over an hour at a variety of enzyme concentrations (Figure 2C). We chose 1.75 μM of enzyme and 20 μM of substrate to use in the high-throughput screen.

### The high-throughput screening for over 5000 chemicals

The screen to identify small molecule inhibitors was performed using an existing custom library of over 5000 compounds aliquoted at 2 different concentrations,1.25 μM or 3.75 μM, in a total of 48 384-well plates. The plates were organised with columns 3-22 containing the compounds; the remaining columns were used for reaction controls including substrate only and substrate with enzyme but no drug (Figure 3Aii). The PLpro enzyme was pre-incubated with the drugs for 10 minutes, and then the reaction was initiated by the addition of substrate. The activity was recorded from 0 to 20 minutes with 3 minutes intervals between readings, this resulted in 7 timepoints which were then used to calculate the slope (Figure 3Ai).

**Figure 3.**
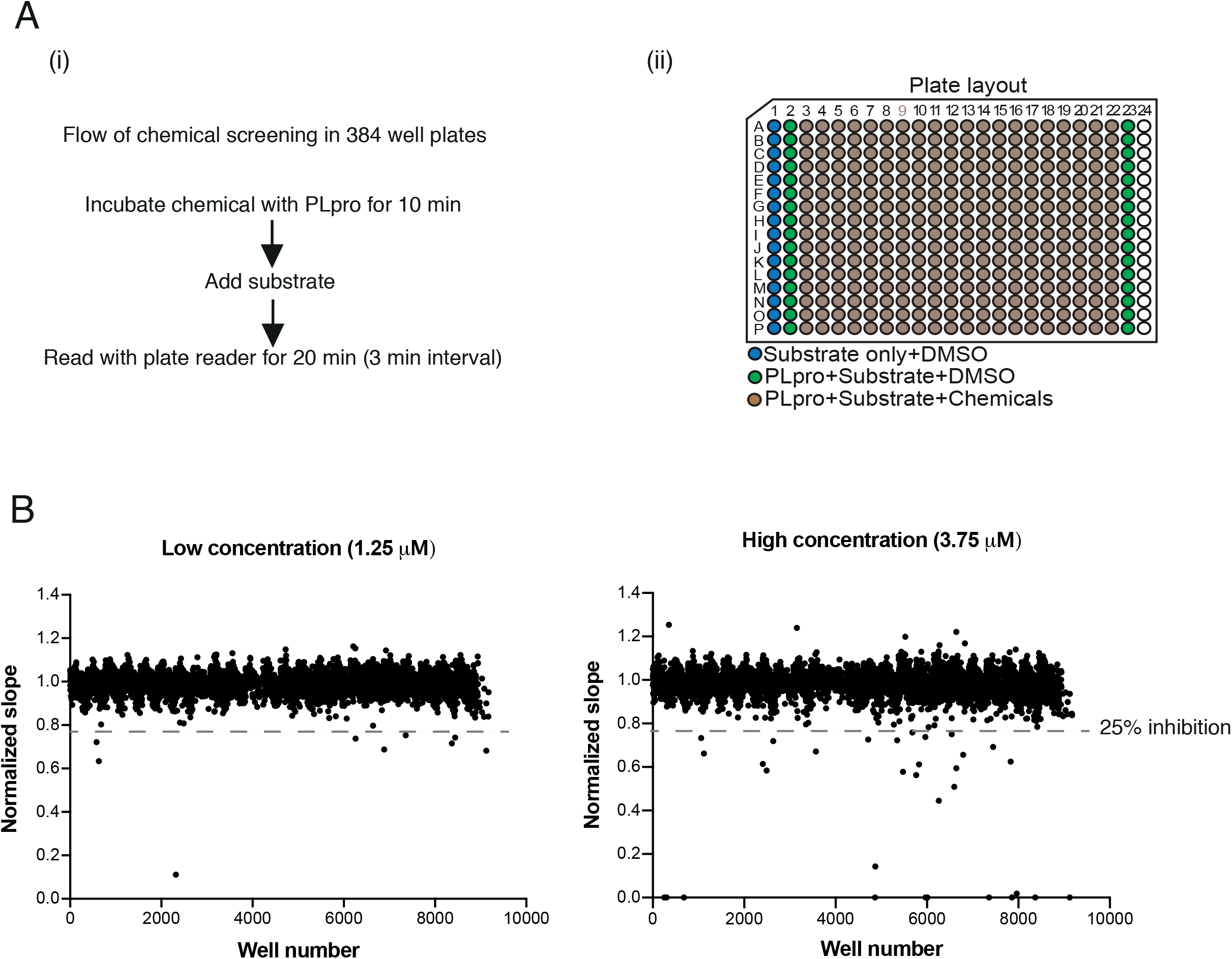
High-throughput screening of the drug library. A. Flow diagram of the drug screen. Over 5000 chemicals were dispensed into 24 of 384-well plates, enzyme was pre-incubated with drug before the addition of substrate and fluorescent intensity was read at an excitation 330 nm and emission wavelength at 420 nm with a Tecan Spark plate reader. B. Results of the screen. Two concentrations of the drug library were aliquoted-1.25 μM and 3.75 μM. A single dot from the scatter plot represents each of the over 5000 chemicals. There are 8 drugs and 29 drugs, in low and high concentration respectively that reduce the PLpro activity equal or more than 25%. Amongst those hits, 4 drugs were overlapping. In the low concentration (1.25 μM), 1 drug is auto-fluorescent and 3 have a Z score of more than −3.5.

We selected a total of 29 candidates from the plates using 3.75 μM drug concentration and a further 8 candidates were selected from the plates with 1.25 μM concentration (Figure 3B) that exhibited apparent inhibition greater than or equal to 25%; 4 of the drugs were overlapping in both concentrations. Amongst the hits, 22 out of 29 drugs from the high concentration plates and 1 drug from the low concentration plates were auto-fluorescent at 420 nm, and were excluded from our list. Three additional candidates were eliminated because the degree of inhibition was weak and the same drugs did not inhibit at higher concentration. This narrowed the final list down to 7 drugs (Table 1).

**Table 1.** Percentage of inhibition and Z score on hits from screening

|  |  | Inhibition (%) |  |  |
| --- | --- | --- | --- | --- |
| No. | Name | 3.75 µM (slope value/control) | 1.25 µM (slope value/control) | Z score |
| 1 | PDK1/Akt/Flt Dual Pathway Inhibitor | 100 (0/491) | 89 (50/452) | -14.53 |
| 2 | Ursodiol | 55 (187/420) | 26 (324/440) | -7.78 |
| 3 | Pyrocatechuic acid | 49 (226/444) | 0 (453/441) | -6.846 |
| 4 | Ro 08-2750 | 42 (287/491) | 19 (366/452) | -5.79 |
| 5 | Cdk4 Inhibitor III | 39 (302/491) | 19 (367/452) | -5.34 |
| 6 | beta-Lapachone | 34 (322/486) | 7 (420/453) | -4.66 |
| 7 | Dihydrotanshinone I | 33 (329/490) | 0 (449/440) | -4.52 |

### Gel-based PLpro assay to test candidates

We wanted to exclude artifacts that might result from the fluorescent-based assay. For this reason we designed a gel-based PLpro protease assay (Figure 4A and B). We constructed a new version of the PLpro substrate that had the 10 amino acids at the junction of nsp2-3 containing the cleavage site attached to GST at the N-terminus and MBP at its C-terminus, resulting in a 67 kDa peptide (Supplementary Figure S3A). If PLpro mediated cleavage occurs, products of 25 kDa and 42 kDa will be generated which can easily be detected and visualized by SDS-PAGE (see Figure 4Ci, lane 3).

**Figure 4.**
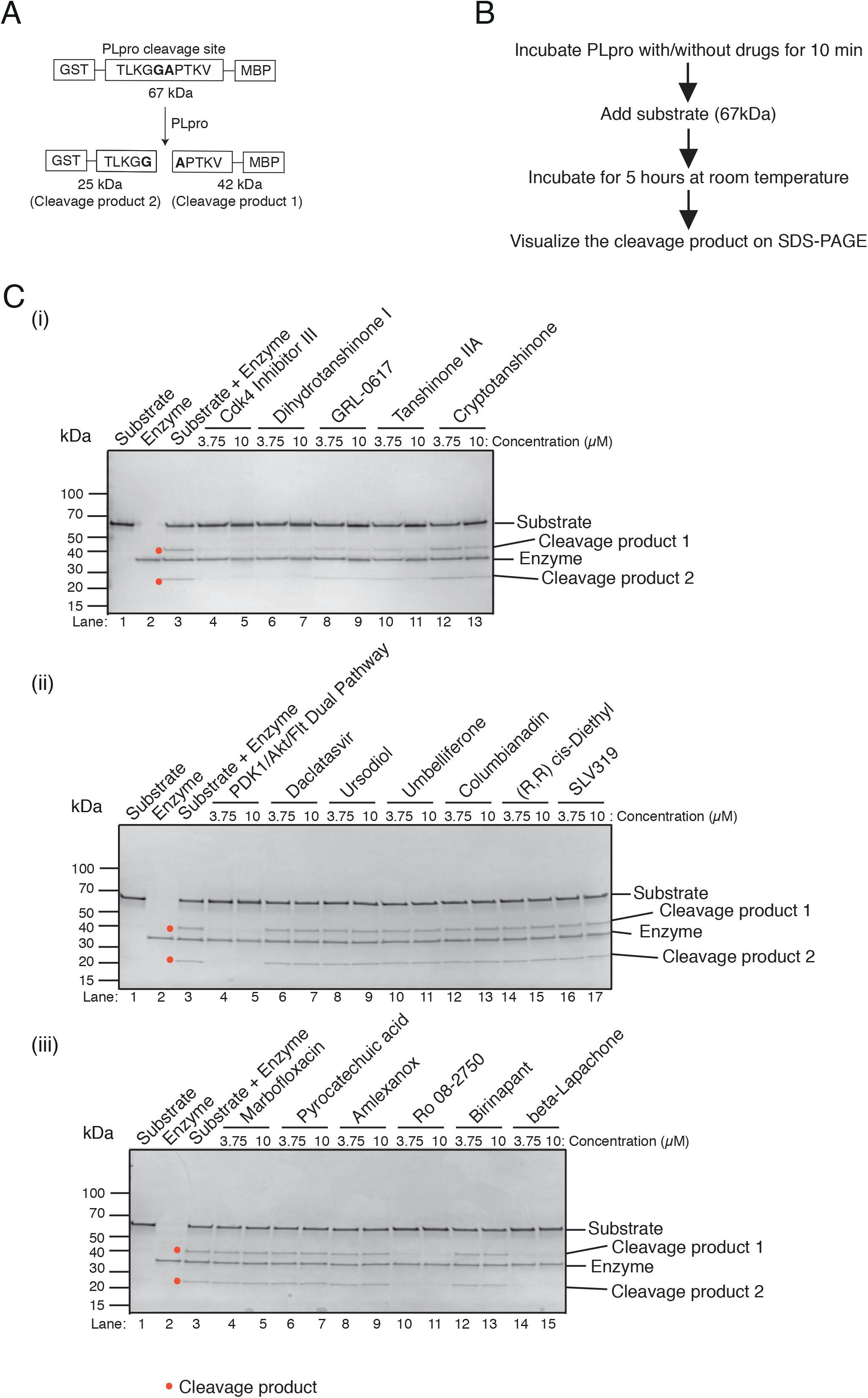
Validation of the hits with a gel-based protease assay. A. Schematic of the substrate designed for the gel-based protease assay. A polypeptide with a N terminal GST and a C terminal MBP domain separated by 10 amino acids from the natural nsp2/3 junction (similar to that used in the fluorescence assay) was constructed. Cleavage between the amino acids G and A results in 25 kDa and 42 kDa products. B. Flow diagram of the gel-based assay. C. (i-iii) Two concentrations of each drug were used, 3.75 μM and 10 μM. Only Cdk4 Inhibitor III, Dihydrotanshinone I, GRL-0617, Tanshinone IIA, Cryptotanshinone, PDK1/Akt/Flt Dual Pathway Inhibitor, Ro 08-2750 (2,3,4,10-Tetrahydro-7,10-dimethyl-2,4-dioxobenzo[g]pteridine-8-carboxaldehyde) and beta-Lapachone show inhibition towards PLpro cleavage. GRL-0617, Tanshinone IIA, and Cryptotanshinone are the published drug inhibitors of PLpro.

One of the hits from the screen, Dihydrotanshinone I was previously shown to inhibit the PLpro from SARS-CoV-1 [20]. Two other tanshinone derivatives, Tanshinone IIA and Cryptotanshinone, were shown to be better inhibitors of this enzyme than Dihydrotanshinone I [20]. In separate studies, a non-covalent inhibitor, GRL-0617, has been reported to be effective against SARS-CoV-1/ −2 PLpro by different groups [21–23]. We, therefore, decided to include these drugs in our validation experiments. As shown in Figure 4C, 5 of 7 candidate hits —PDK1/Akt/Flt Dual Pathway Inhibitor (Figure 4Cii), Cdk4 Inhibitor III (Figure 4Ci), Dihydrotanshinone I, Ro 08-2750 (Figure 4Ciii) and beta-Lapachone — strongly inhibited the enzyme activity at both drug concentrations tested as indicated by the reduced levels of the 25 and 42 kDa products (red dot in Figure 4). Two of the final 7 drugs, Ursodiol and Pyrocatechuic acid, did not inhibit PLpro cleavage of the substrate in this assay and, therefore, appear to be false positives. We tested some of the auto-fluorescent hits, but none inhibited cleavage of the peptide, further confirming them as false positives (Figure 4Cii and iii). Tanshinone IIA and Cryptotanshinone also inhibited cleavage, but only at the higher concentration. Thus, Dihydrotanshinone I appears to be the strongest inhibitor amongst the Tanshinone derivatives. Similarly, GRL-0617 also inhibited cleavage only at the highest concentration. 3CLpro, contained in the nsp5 protein, is the other main protease encoded by SARS-CoV-2, and, like PLpro, 3CLpro is also has an active site cysteine. However, none of the five screen hits inhibited 3CLpro activity in an analogous gel-based assay (Milligan, Zeisner et al. this series; Supplementary Figure S3B).

We determined the IC_50_ value for all 5 inhibitors from the screen that were validated by gel-based assay. All of them are below 1 μM: 0.26 μM for PDK/Akt/Flt Dual Pathway Inhibitor, 0.53 μM for Ro 08-2750, 0.39 μM for Cdk4 Inhibitor III, 0.61 μM for beta-Lapachone, and 0.59 μM for Dihydrotanshione I (Figure 5A-E). The IC_50_ for the 3 published drugs was slightly higher in our experimental conditions: 1.79 μM for GRL-0617, 1.57 μM for Tanshinone IIA, and 1.34 μM for Cryptotanshinone (Figure 5F-H), consistent with the gel-based assay results. During this time we realised that PDK1/Akt/Flt Dual Pathway Inhibitor was also identified in several other screens being performed in the lab. We, therefore, considered it to likely to be non-specific and we eliminated it from further consideration.

**Figure 5.**
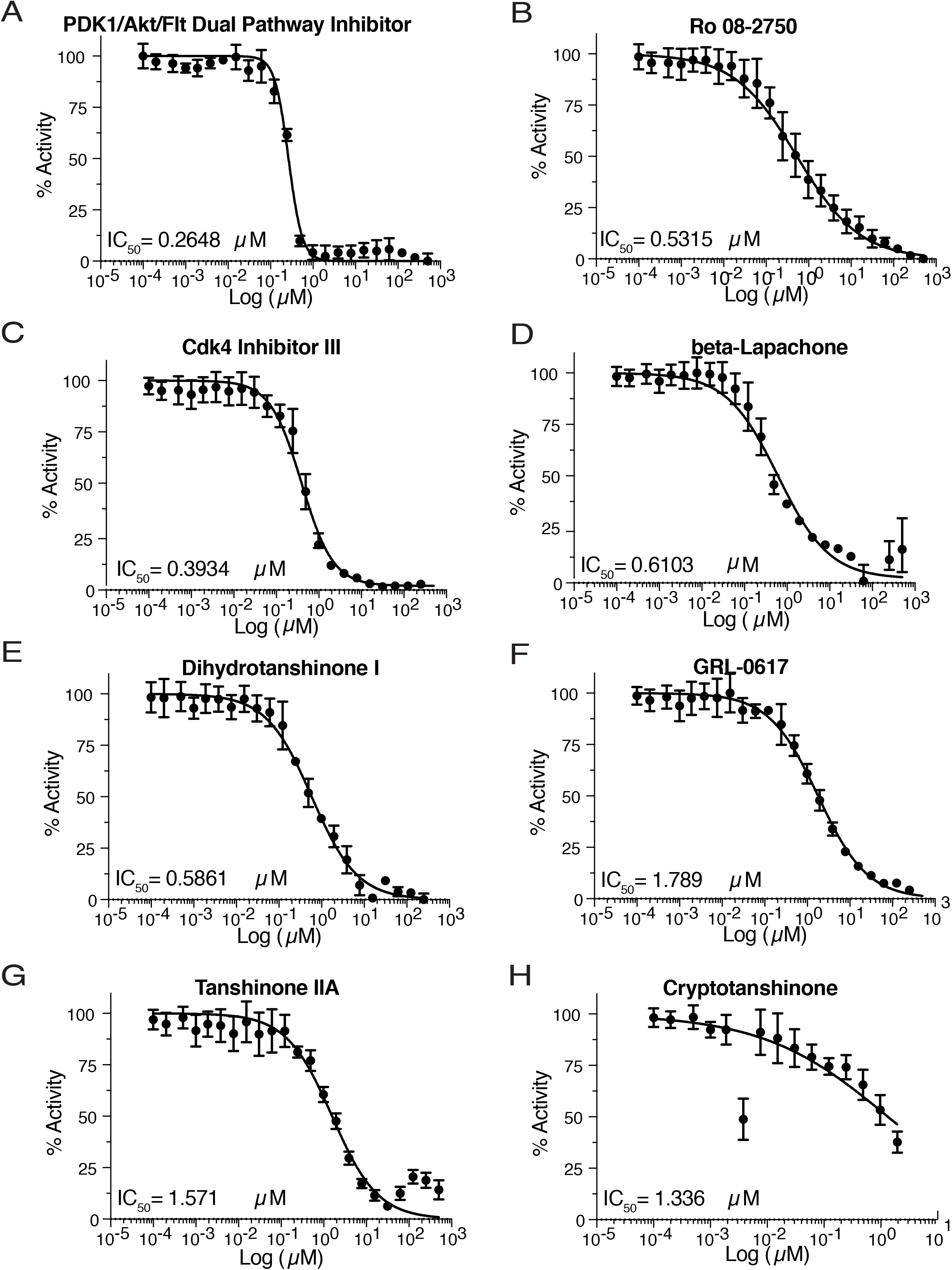
Determination of the IC_50_ of the validated hits with the fluorescent-based assay. A. PDK1/AKT/Flt Dual Pathway Inhibitor, B. Ro 08-2750, C. Cdk4 Inhibitor III, D. beta-Lapachone, E. Dihydrotanshinone I, F. GRL-0617, G. Tanshinone IIA, H. Cryptotanshinone.

### Inhibition of PLpro isopeptidase activity

We performed orthogonal assays using two different substrates of PLpro to test the inhibition of isopeptidase activity of PLpro by the small molecule inhibitors identified in the screen. When K48-linked triubiquitin (Ub3) is incubated with PLpro, it is efficiently cleaved to diubiquitin (Ub2) and monoubiquitin (Ub1) as the final products [18]. Similarly, pro-ISG15 is cleaved to ISG15. When PLpro is pre-incubated with 10 μM of beta-Lapachone or Dihydrotanshinone I, we observed potent inhibition of both K48-linked Ub3 and pro-ISG15 cleavage whereas moderate inhibition was observed for both Ro 08-2750 and the previously identified inhibitor of PLpro, GRL-0617, at these concentrations (Figure 6). A potential 3CLpro inhibitor, shikonin, did not inhibit isopeptidase activity with either substrate. These experiments demonstrate the inhibitory effect of these small molecules across a spectrum of PLpro substrates.

**Figure 6.**
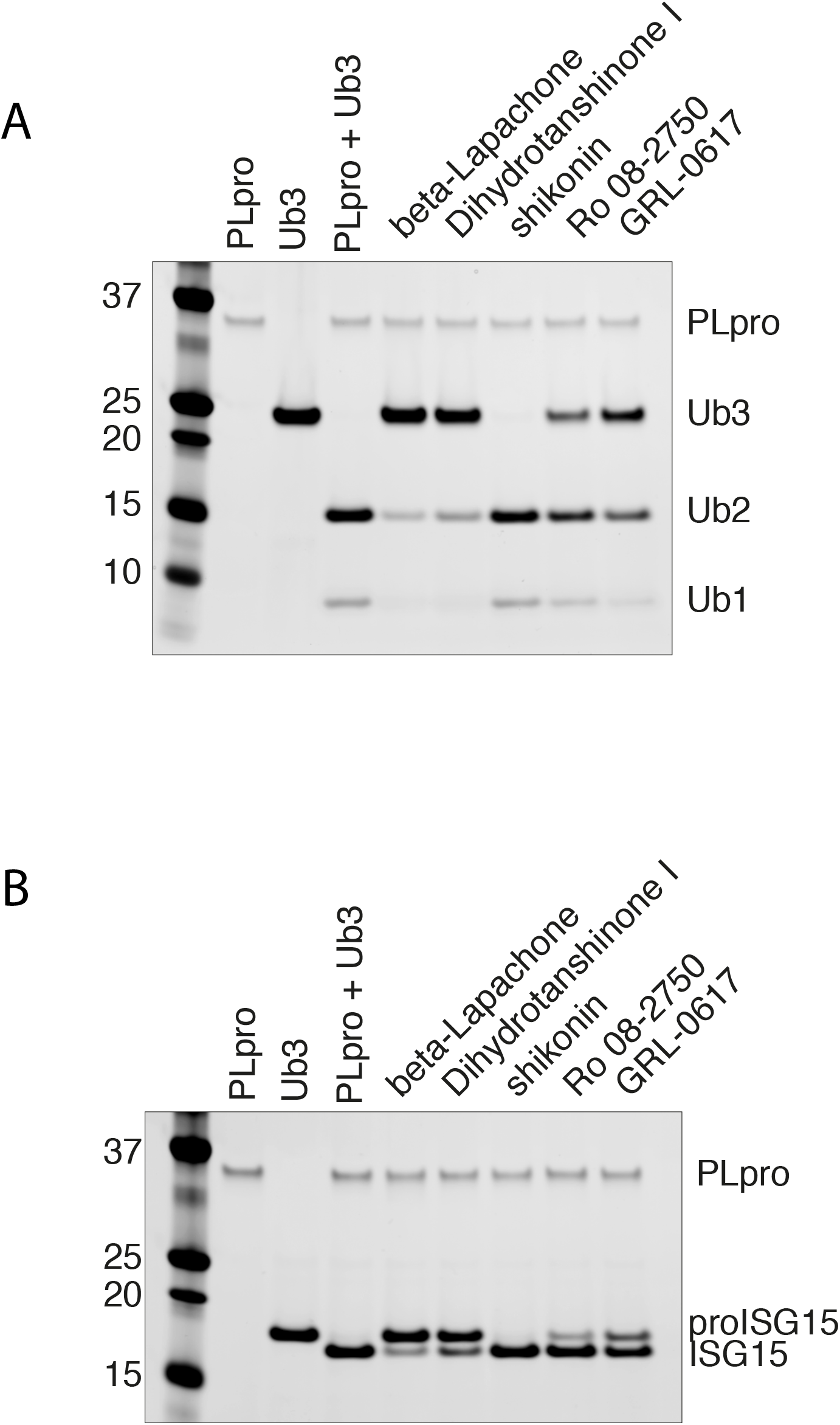
Inhibition of PLpro isopeptidase activities. A. PLpro DUB activities. PLpro cleaves K48-linked Ub3 to Ub2 and Ub1. These activities were inhibited strongly by beta-Lapachone and Dihydrotanshinone I, moderately by Ro 08-2750 and GRL-0617, but not by shikonin. B. PLpro deISGylating activities. Pro-ISG15 is being cleaved to ISG15 by PLpro. Similar observation as in A.

### Cell culture-based antiviral proliferation assay

We next tested the ability of these compounds to inhibit viral growth in a cell culture-based assay where VERO E6 cells are infected with SARS-CoV-2. Two of the drugs, beta-Lapachone and Cdk4 Inhibitor III, were cytotoxic in the low micromolar range and were not pursued (Supplementary Figure S4A and B). The two tanshinone derivatives not identified in our screen (Cryptotanshinone and Tanshinone IIA) did not inhibit viral growth below 200 μM (Supplementary Figure S4C and D). Ro 08-2750 and GRL-0617 were better at inhibiting viral growth (EC_50_ 20 and 32.6 μM respectively) but also exhibited some cell toxicity at higher concentrations (Figure 7Bi, Ci and Supplementary Figure S5). Dihydrotanshinone I proved to be the best inhibitor since it effectively inhibited the SARS-CoV-2 proliferation at an EC_50_ of 8 μM (Figure 7Ai) and did not exhibit much cytotoxicity, even at high concentrations.

**Figure 7.**
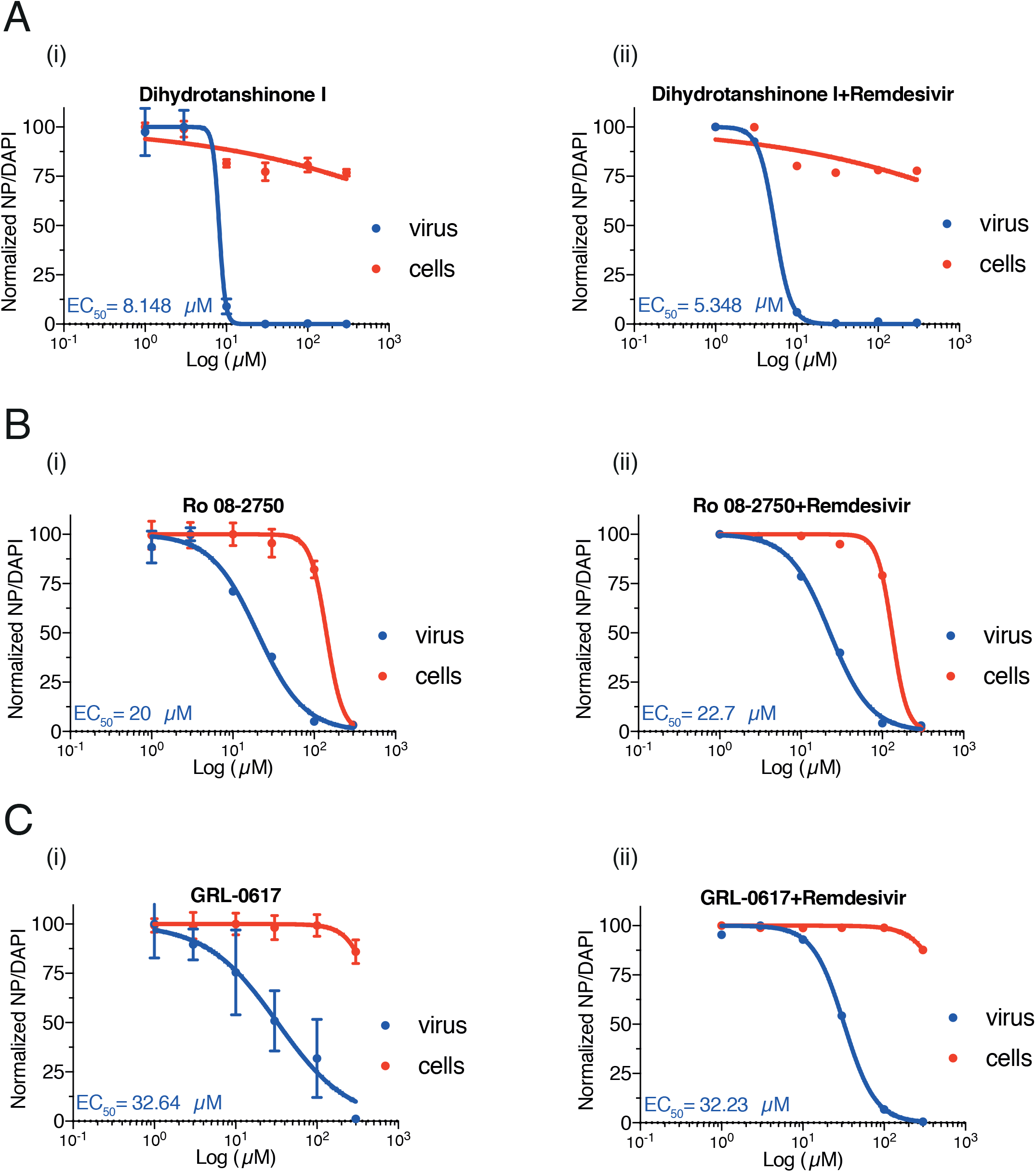
Determination of the EC_50_ of Dihydrotanshinone I (Ai), Ro 08-2750 (Bi), GRL-0617 (Ci) in a cell culture-based assay and the synergistic effect of each drug with remdesivir (Aii, Bii and Cii). Data was collected from three replicates.

Since remdesivir is the one and only approved antiviral compound for COVID-19, we wanted to test whether any of the drugs we have found show any synergy with remdesivir (Supplementary Figure S5). However, addition of 0.5 μM remdesivir, a concentration just below that required to inhibit viral growth, did not reduce the EC_50_ of any of our compounds (Figure 7A-Cii).

## Discussion

Dihydrotanshinone I, which emerged as the best overall hit from our screen, is a natural compound isolated from lipophilic fraction of *Salvia miltiorrhiza*, which has a long history in traditional Chinese medicine [24]. Several derivatives of Tanshinones were previously reported to be inhibitors for SARS-CoV-1 PLpro and to a lesser extent in 3CLpro [20, 24]. We have found that Dihydrotanshinone I is the best inhibitor of SARS-CoV-2 PLpro and did not inhibit 3CLpro. Although we, like other groups [23, 25, 26], used a truncated nsp3 protein including just the Ubl2 and PLpro domains, and there is evidence that cleavage by the full length nsp3 may have slightly different specificities [18]. The fact that Dihydrotanshinone I stops viral replication suggests that it is a good nsp3 PLpro inhibitor in cells.

PLpro also recognizes and removes K48-linked polyubiquitin chains (Ub) and ISG15 from host cell target proteins. It is known that either ubiquitin or ISG15 are covalently bonded to target proteins during the cellular response to viral infection. The deubiquitinating (DUB) and deISGylating activities of PLpro following LXGG motifs thus have an implication in viral invasion by shutting down the viral induced host innate immune response [27]. PLpro of SARS-CoV-2 further shows preferential activity in cleaving ISG15 over Ub in comparison to SARS-CoV-1 [18, 21, 23]. Interestingly, although the high transmission (and potentially more deadly) U.K. variant of SARS-CoV-2, B.1.1.7 was believed attributed to multiple mutations in the spike protein, the significance of a particular point mutation, A1708D, in PLpro for viral invasion remains unexplored [28].

## Experimental Procedures

### Expression constructs

The coding sequence of SARS-CoV-2 nsp3 1564-1878 amino acid (NCBI reference sequence NC_045512.2) was selected as previously reported [14]. The His-TEV bacterial sequence was codon optimized (Table S2) and cloned into plasmid pET11a at NdeI/BamHI sites (synthesised and cloned by GeneWiz) (Addgene ID 169192). The His-Sumo bacterial version was codon optimized and cloned into K27-Sumo (Addgene ID 169193) via NEBuilder HiFi DNA Assembly Cloning Kit (NEB). Vector template was amplified using primers oEcoli-C_48 and 49, PLpro sequence was amplified from His-TEV bacteria strain using primers oEcoli-C_51 and 52.

### Expression and purification

#### Bacteria

His-TEV and His-Sumo constructs were introduced into T7 express lysY/I^q^ *E. coli* cell (NEB) for expression. Cells were grown at 37°C to log phase to achieve OD 0.8. Cells were then induced by the addition of 0.5 mM IPTG and switched to 18°C to incubate overnight. Cells were harvested and lysed in buffer A (50 mM Tris-HCl, pH 7.5, 10% glycerol, 1 mM DTT, 0.02% NP-40, 500 mM NaCl and 30 mM imidazole), with the addition of 100 ug/ml lysozyme and sonicated 24 × 5 sec. Lysates were centrifuged and the supernatant was collected. The supernatant was incubated with Ni-NTA agarose beads (Thermo) for 2 hrs at 4°C. Beads were washed with wash buffer A. The protein was eluted with 200 mM (His-TEV) or 400 mM (His-Sumo) of imidazole. Fractions were pooled and dialyzed in buffer B (50 mM Tris-HCl, pH 7.5, 10% glycerol, 1 mM DTT, 0.02% NP-40 and 50 mM NaCl) and 0.1 mg/ml His-TEV protease (His-TEV) or 0.02 mg/ml His-Ulp1 (His-Sumo) to cleave-off the His-TEV- and His-Sumo-tag respectively. After dialysis the lysate was incubated with Ni-NTA agarose beads once again to remove the proteases. The flow through was collected and loaded onto a MonoQ 5/50 GL column (GE healthcare) with buffer B, with gradient from 0.1 M to 1 M NaCl. Flow through was collected and concentrated using Amicon ultra 10 kDa (Merck). It was then loaded onto a Superdex S200 Increase 10/300 GL (GE healthcare) with buffer C (25 mM HEPES-KOH, pH 7.6, 10% glycerol, 0.02% NP-40, 150 mM NaCl and 2 mM DTT). Peak fractions were collected and pooled.

#### Baculovirus

3xFlag-His_6_-PLpro (3FH-PLpro) was expressed in baculovirus-infected insect cells. The coding sequence was codon-optimised for *S. frugiperda* and synthesised (GeneArt, Thermo Fisher Scientific). PLpro DNA was subcloned into the biGBac vector pLIB [29] to include an N-terminal 3xFlag-His_6_ tag (sequence: MDYKDHDGDYKDHDIDYKDDDDKGSHHHHHHSAVLQ) (Addgene ID 169194). Baculoviruses were generated using the EMBacY baculoviral genome [30] in Sf9 cells (Thermo Fisher Scientific). For protein expression Sf9 cells were infected with baculovirus and collected 48 hours after infection, flash-frozen and stored at −70°C.

### PLpro gel-based assay substrate

TLKGG//APTKV (nsp2/3 junction) was used as the substrate for gel-based assay. GST-tag and MBP-tag were attached to its N- and C-terminus respectively (Addgene ID 169195). The cleavage at G//A resulted in 25 and 42 kDa products. The constructs were expressed in T7 express lysY/I^q^ *E. coli* cell (NEB) as follows. Cells were grown at 37°C to log phase to achieve OD 0.8. Cells were then induced by the addition of 1 mM IPTG at 37°C for 4 hrs. Cells were harvested and lysed in buffer D (50 mM Tris-HCl, pH 7.5, 10% glycerol, 1 mM DTT, 0.02% NP-40, 300 mM NaCl), with the addition of 1X protease inhibitor Leupeptin (Merck), Pepstatin A (Sigma), AEBSF (Sigma), 100 ug/ml lysozyme and sonicated for 24 × 5 sec. Lysate was centrifuged and supernatant was collected. The supernatant was incubated with amylose resin (NEB) for 2 hrs at 4°C. Beads were washed with wash buffer D. The protein was eluted with 10 mM maltose. Peak fractions were collected and concentrated using Amicon ultra 10 kDa (Merck). It was then loaded on Superdex S200 Increase 10/300 GL (GE healthcare) with buffer C. Peak fractions were collected and pooled.

### Ulp1 catalytic fragment

An expression plasmid pFGET19-Ulp1 from Addgene. Plasmids were transformed into T7 express lysY/I^q^ *E. coli* cell (NEB) for expression. Cells were grown at 37°C to log phase to achieve OD 0.8. Cells were then induced by the addition of 1 mM IPTG at 37°C for 4 hrs. Cells were harvested and lysed with in buffer E (50 mM Tris-HCl, pH 8, 5 mM magnesium acetate, 10% glycerol, 1 mM DTT, 0.02% NP-40, 500 mM NaCl, 20 mM imidazole), with the addition of 1X protease inhibitor Leupeptin (Merck), Pepstatin A (Sigma-Aldrich), AEBSF (Sigma-Aldrich), 100 ug/ml lysozyme and sonicate 24×5s. Lysate was centrifuged and supernatant was collected. The supernatant was incubated with Ni-NTA agarose beads (Thermo Scientific) for 1 hr at 4°C. Beads were washed with wash buffer D. The protein was eluted with 250 mM imidazole. Peak fractions were collected and concentrated using Amicon ultra 10 kDa (Merck). It was then loaded on Superdex S200 Increase 10/300 GL (GE healthcare) with buffer F (50 mM Tris-HCl, pH 8, 5 mM magnesium acetate, 10% glycerol, 0.5 mM TCEP, 0.01% NP-40, 500 mM NaCl). Peak fractions were collected and pooled.

### Fluorescent-based assay

Three synthetic peptide substrates were used- Pro1: ELNGG//AYTRY (nsp1-2 junction), Pro2: TLKGG//APTKV (nsp2-3 junction) and Pro3: FTLKGG//APTKVT (nsp2-3 junction). Anthranilate (2-aminobenzoyl-Abz) and nitro-L-tyrosine [Y(3-NO_2_)R] were attached at the N- and C-termini as fluorescent donor and quencher respectively. The substrates were made in-house by the Peptide Synthesis department in the Francis Crick Institute. Fluorescence was read on a Spark multimode microplate reader (Tecan) with an excitation wavelength at 330 nm and emission wavelength at 420 nm. The buffer used for the assay was buffer G: 50 mM HEPES-KOH, pH 7.6, 2 mM DTT, 10% glycerol, 0.02% Tween-20, 20 μM substrate and 1.75 μM enzyme, unless it was stated in the figure.

### Enzymatic assay

Kinetic constants were measured using a fluorescent-based assay. 0.5 μM enzyme were used in the titration of substrate Pro3 from 800 to 1.43 μM with 0.75-fold. Vmax and Km were calculated from non-linear Michaelis-Menten via linear regression of slope value.

### High-throughput inhibitor screening

A total of over 5000 chemicals (Sigma, Selleck, Enzo, Tocris, Calbiochem, and Symansis) were obtained from High-Throughput Screening (HTS) facility in Francis Crick Institute. The drugs were aliquoted into 48 384-well plates in 2 concentrations with final concentration of 1.25 μM and 3.75 μM in 20 μl reaction. 5 μl of enzyme was pre-incubated with drugs for 10 min before the addition of 15 μl substrate.

Fluorescence was monitored at 2 minutes for 20 minutes with 3 minutes intervals giving a total of 7 readings.

### Data analysis

MATLAB was used to process data (more details in nsp13 paper in this issue)

### Gel-based assay

0.5 μM of enzyme was pre-incubated with the selected inhibitor drugs for 10 minutes. 0.5 μM of gel-based assay substrates were added and incubated at RT for 5 hrs. The reactions were ran on 4-15% TGX gel (BioRad, Hercules, CA) and stained with Instant blue stain (Expedeon).

### Assays for deubiquitylation and deISGylation activity

Expression and purification of PLpro and the substrates K48-linked Ub3 and pro-ISG15 were as described previously [18]. PLpro was diluted in buffer G (50 mM HEPES-KOH pH 7.6, 2 mM DTT, 10% (v/v) glycerol, 0.02% (v/v) Tween-20) to 100 nM and pre-incubated with indicated concentrations of inhibitor at room temperature for 10 mins. The substrates Ub3 and pro-ISG15 were diluted in buffer G to a concentration of 2 μM. An equal volume of substrate was then mixed with the inhibited PLpro to give a final concentration of 50 nM PLpro, 10 μM inhibitor and 1 μM substrate. The reaction was incubated for 1 hour at 25°C and the reactions were stopped by addition of LDS sample buffer. The samples were separated on 4-12% NuPAGE Bis-Tris Gels (ThermoFisher Scientific) and stained using Pierce Silver Stain Kit (ThermoFisher Scientific).

### Cell-based assay

SARS-CoV-2 production, infection and recombinant mAb production was done as described by Zeng, Weissmann, Bertolin et al. this series.

### Data Availability Statement

All data in this paper can be found in FigShare (https://figshare.com/).

## Author Contributions

**Chew Theng Lim**: Conceptualisation, Methodology, Validation, Formal analysis, Investigation, Resources, Writing – Original Draft, Writing – Review and Editing, Visualisation. **Kang Wei Tan**: Conceptualisation, Methodology, Validation, Formal analysis, Investigation, Resources, Writing – Original Draft, Writing – Review and Editing, Visualisation. **Mary Wu**: Methodology, Investigation, Resources. **Rachel Ulferts**: Methodology, Investigation. **Lee A. Armstrong**: Investigation, Writing – Original Draft, Visualisation. **Eiko Ozono**: Investigation, Validation. **Lucy S. Drury:** Investigation, Validation, Writing – Original Draft, Visualisation. **Jennifer C. Milligan**: Investigation, Methodology. **Theresa U. Zeisner:** Methodology. **Jingkun Zeng**: Software. **Florian Weissmann**: Resources. **Berta Canal**: Resources. **Ganka Bineva-Todd**: Resources. **Nicola O’Reilly**: Resources. **Micheal Howell**: Supervision. **Rupert Beale**: Supervision. **Yogesh Kulathu**: Supervision. **Karim Labib**: Supervision. **John F.X Diffley**: Conceptualisation, Methodology, Writing – Review and Editing, Supervision, Project administration, Funding acquisition.

## Acknowledgements

We thank Anne Early and Agustina P. Bertolin for their assistance and High-Throughput Screening (HTS) for dispensing the compound libraries. Theresa U. Zeisner received funding from the Boehringer Ingelheim Fonds. Berta Canal and Florian Weissmann received funding from the European Union’s Horizon 2020 research and innovation programme under the Marie Skłodowska-Curie grant agreement Nos 895786 and 844211 respectively. Yogesh Kulathu and Lee A. Armstrong are supported by Medical Research Council UK (MC_UU_00018/3), EMBO Young Investigator Programme (Yogesh Kulathu), ERC Starting grant (677623) (Yogesh Kulathu) and Lister research prize (Yogesh Kulathu). This work was supported by the Francis Crick Institute, which receives its core funding from Cancer Research UK (FC001066), the UK Medical Research Council (FC001066), and the Wellcome Trust (FC001066). This work was also funded by a Wellcome Trust Senior Investigator Award (106252/Z/14/Z) to J.F.X.D.

## Supplemental Materials

**Figure S1.**
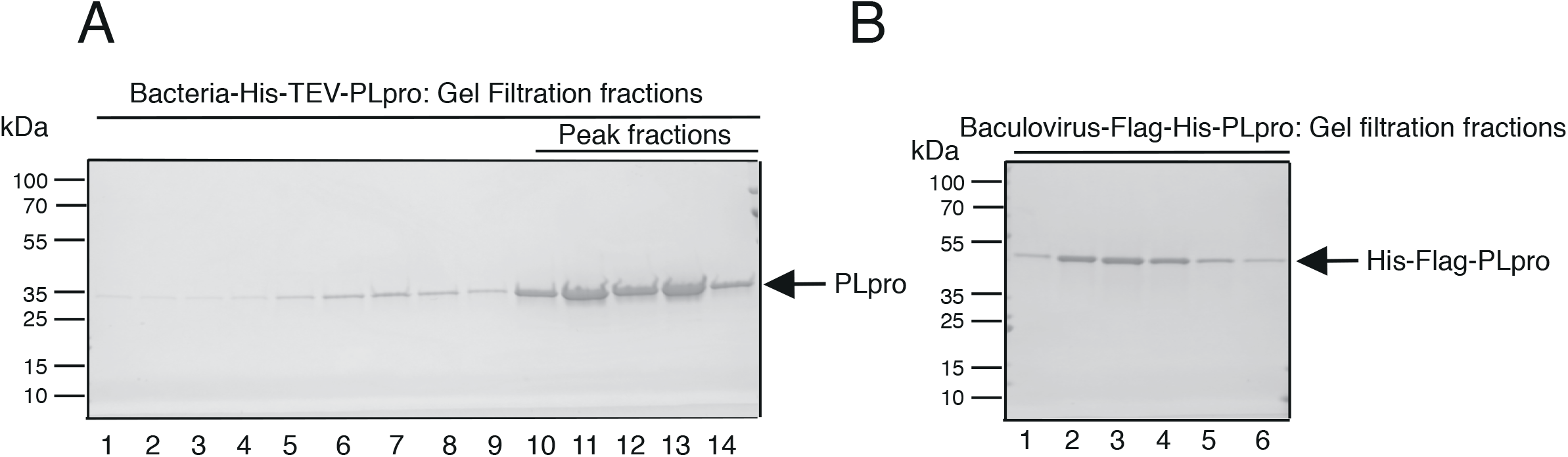
(Related to Figure 1) Purification of PLpro in bacteria and baculovirus. A. Purification of His-TEV-PLpro from bacteria. Lanes 11-13 contain monomer of PLpro from the last step of the purification on a gel filtration column and these were pooled B. Purification of Flag-His-PLpro from baculovirus. Lane 2-4 contain monomer of PLpro from a gel filtration column and were pooled. The tag remains intact with protein.

**Figure S2.**
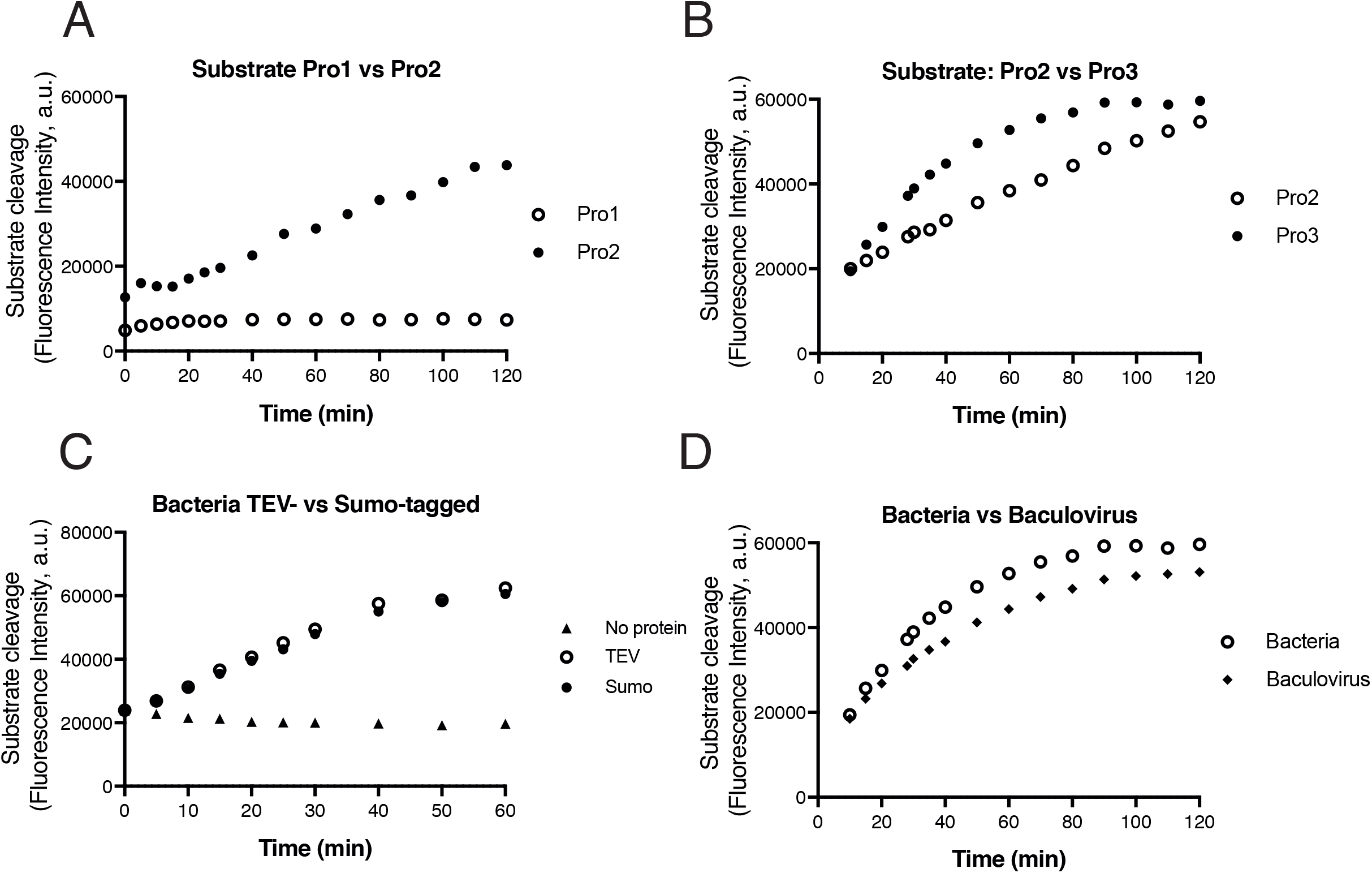
(Related to Figure 2) Optimalization of the FRET assay for PLpro. A. PLpro was able to cleave substrate Pro2 (nsp2/3) but not Pro1 (nsp1/2) (see Material & Methods) B. PLpro was able to cleave substrate Pro3 (12 amino acids) more efficiently than Pro2 (10 amino acids). C. Comparison between the TEV-tagged and Sumo-tagged versions of PLpro enzyme activities. D. Comparison between the baculovirus and bacterial (TEV-tagged) purified PLpro.

**Figure S3.**
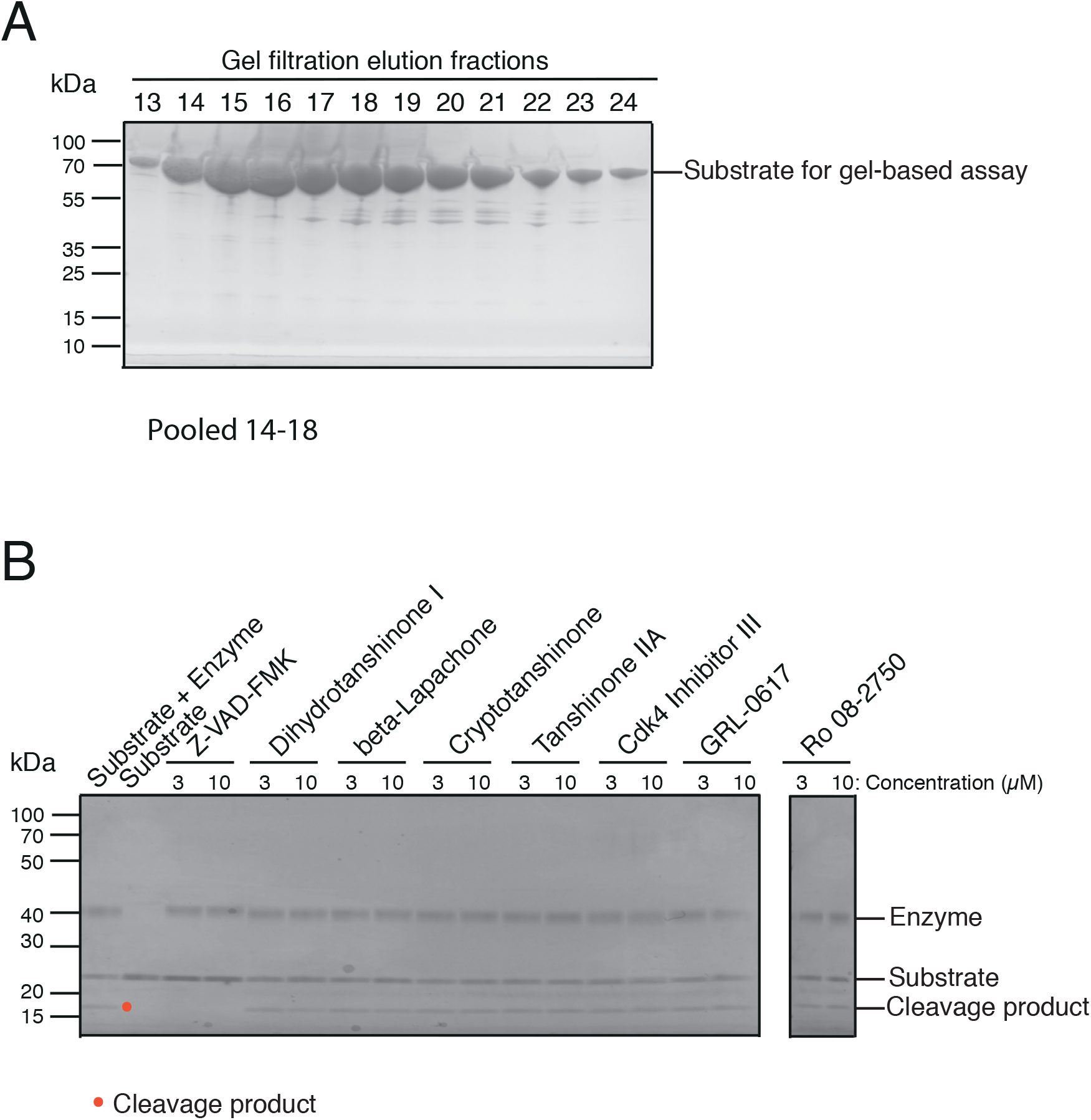
(Related to Figure 4) Gel-based assay. A. Purification of the substrate (67 kDa) with Superdex S200. Fractions 14-18 were pooled and used for the gel-based assay. B. Dihydrotanshione I, beta-Lapachone, Cryptotanshinone, Tanshinone IIA, Cdk4 Inhibitor III, GLR-0617, and Ro 08-2750 did not inhibit 3CLpro (nsp5) activity. Z-VAD-FMK is a novel inhibitor discovered against nsp5 (reference nsp5 paper)

**Figure S4.**
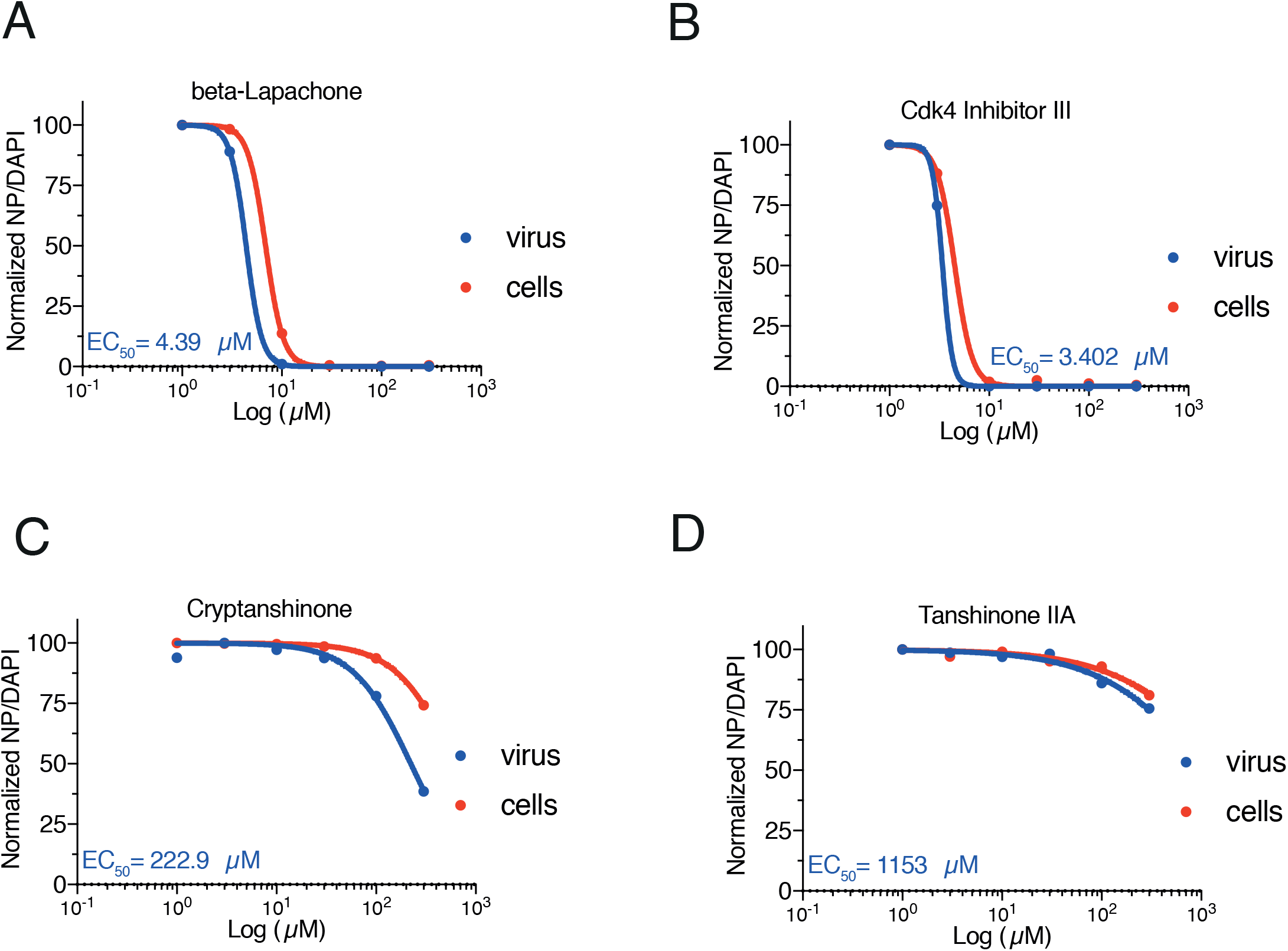
EC_50_ of beta-Lapachone (A), Cdk4 Inhibitor III (B), Cryptotanshinone (C), and Tanshinone IIA (D) from the cell culture-based viral proliferation assay.

**Figure S5.**
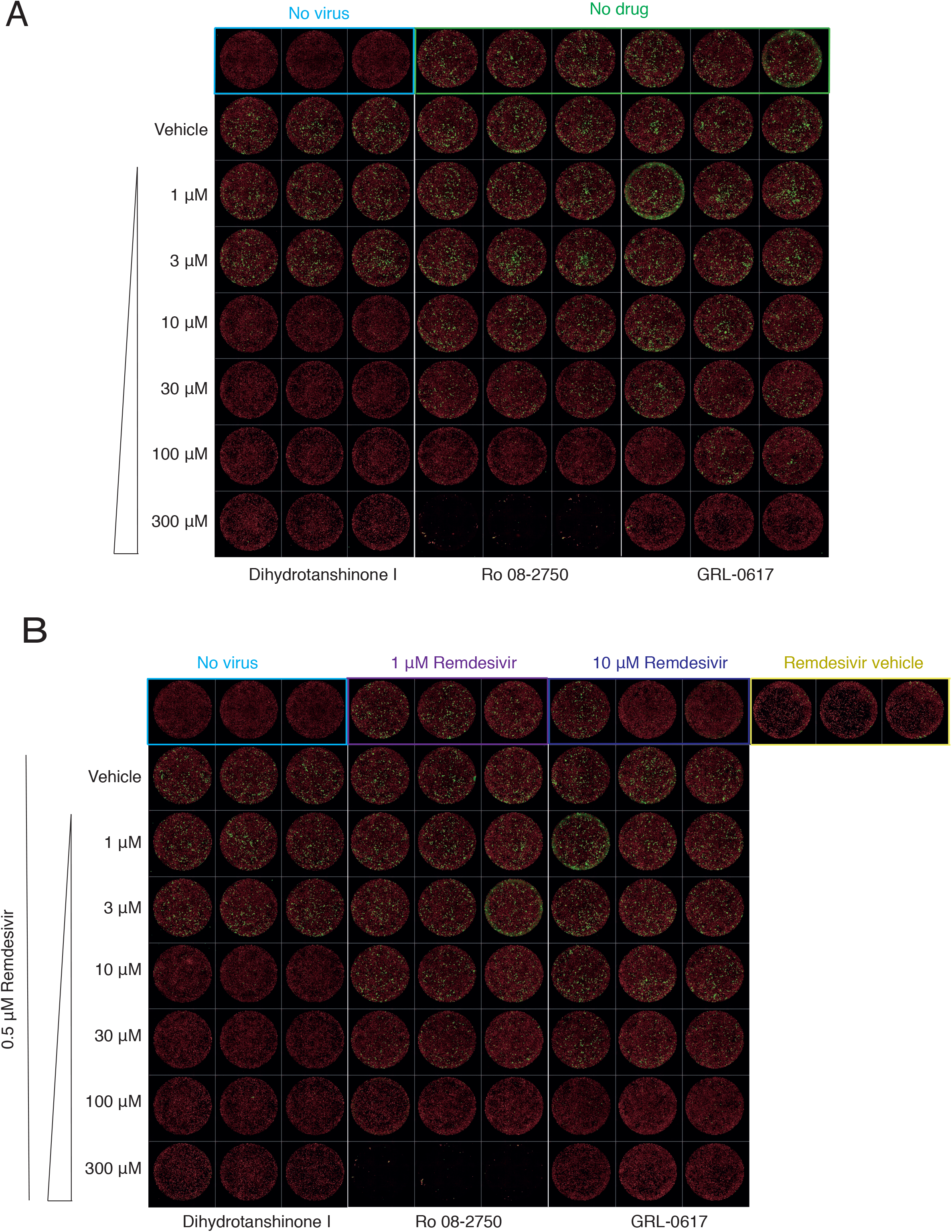
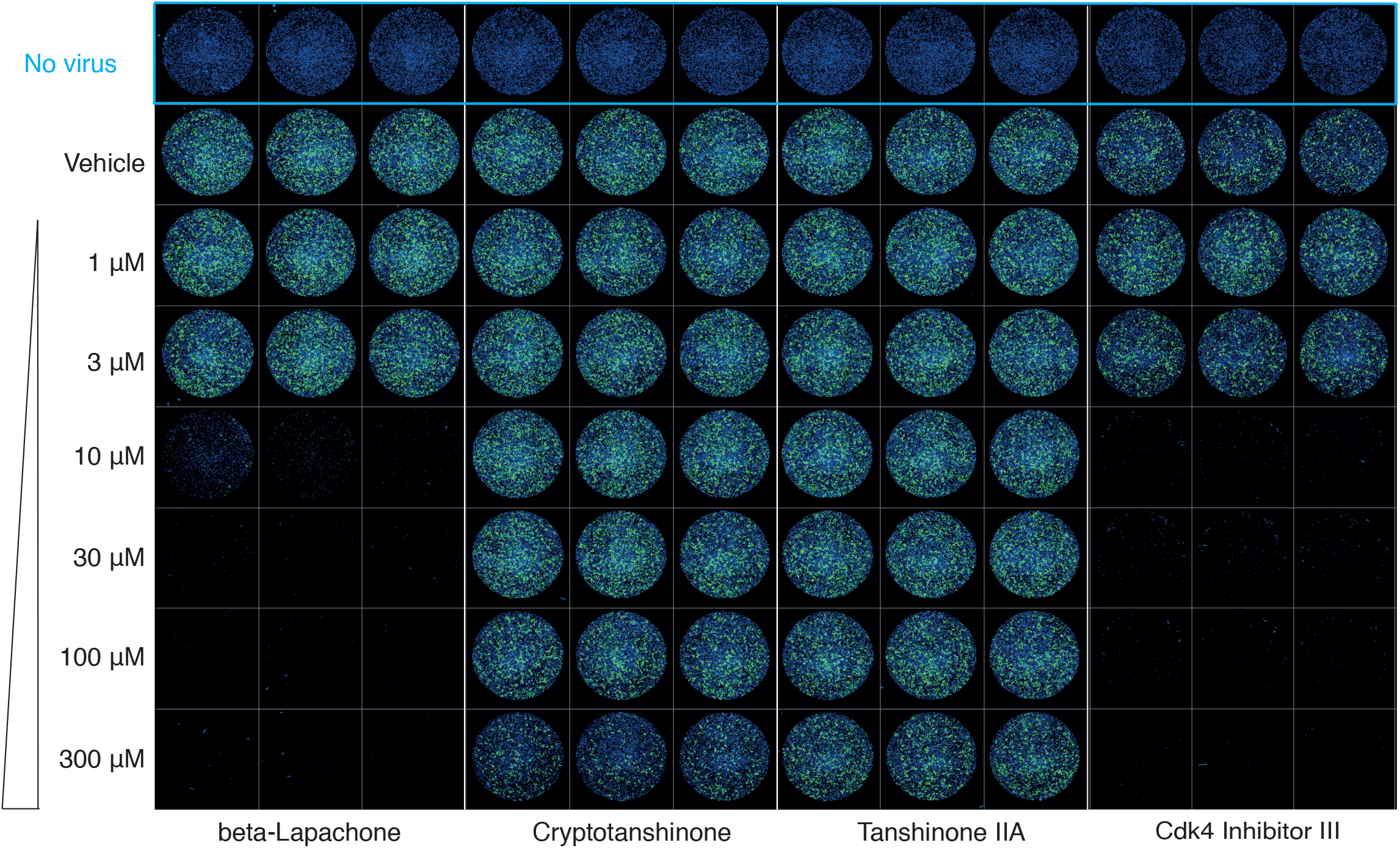
(Related to Figure 7) (A and C) Viral proliferation assay for Dihydrotanshinone I, Ro 08-2750, GRL-0617, beta-Lapachone, Cryptotanshinone, Tanshinone IIA, and Cdk4 Inhibitor III. B. There is no obvious additional effect of combining remdesivir with drugs in (A). These drugs either do not stop COVID-19 viral proliferation or are cytotoxic.

**Table S1.** Chemical list of over 5000 chemicals with scores from low and high concentration.

**Table S2.**
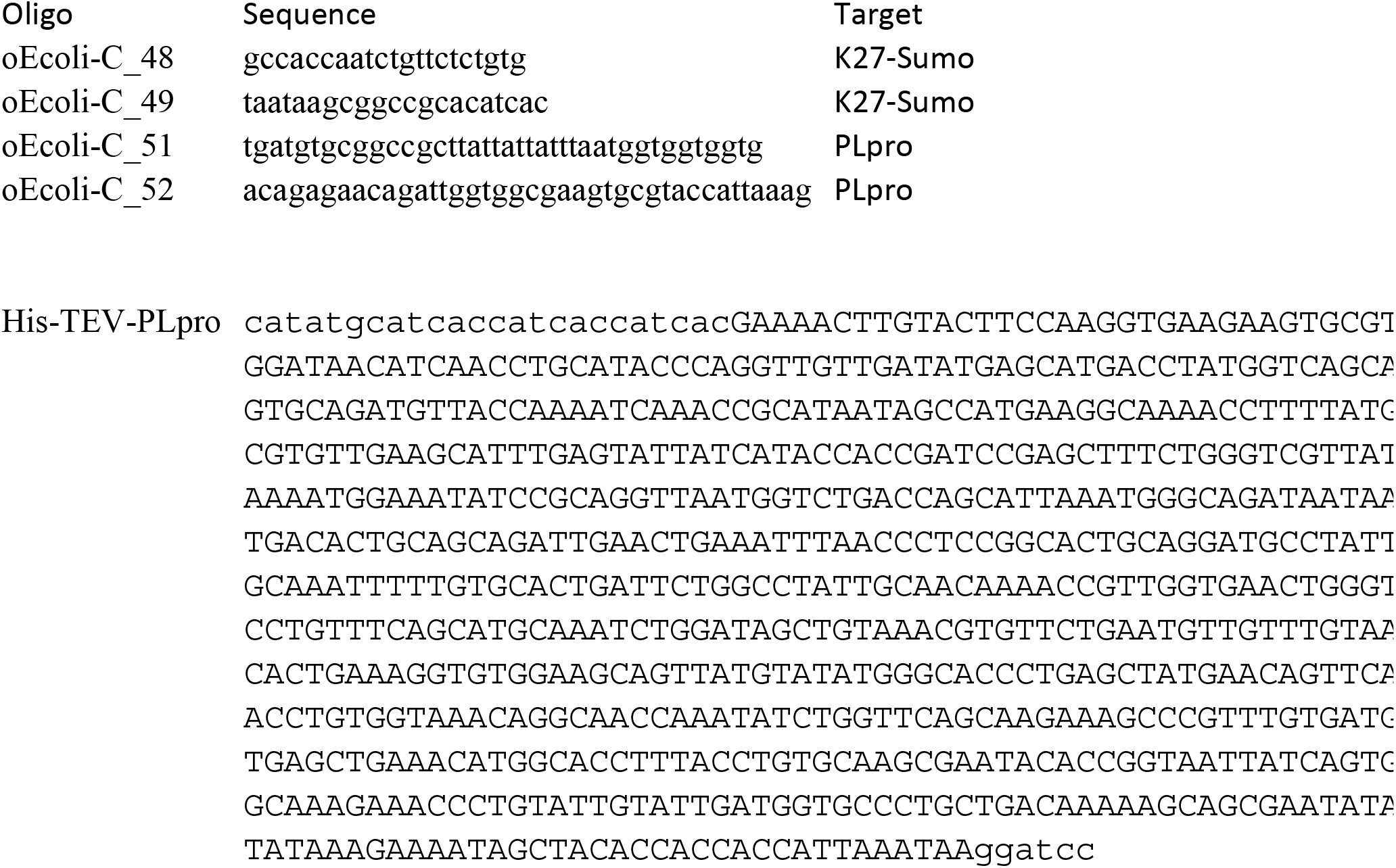
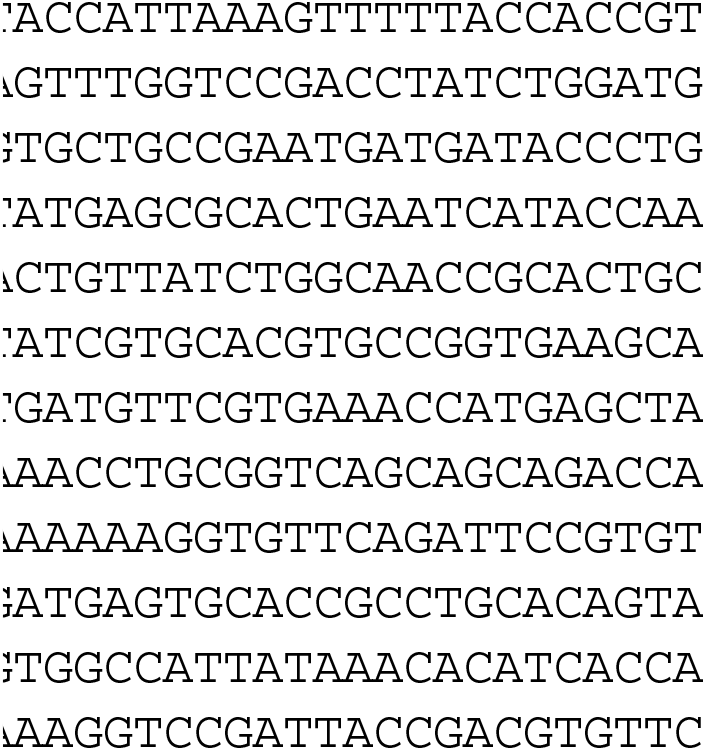
Oligomers and PLpro optimized sequence.

